# Arginyltransferase1 drives a mitochondria-dependent program to induce cell death

**DOI:** 10.1101/2024.11.22.624728

**Authors:** Akhilesh Kumar, Corin R. O’Shea, Vikas K. Yadav, Ganapathi Kandasamy, Balaji T. Moorthy, Evan A. Ambrose, Aliya Mulati, Flavia Fontanesi, Fangliang Zhang

## Abstract

Cell death regulation is essential for stress adaptation and/or signal response. Past studies have shown that eukaryotic cell death is mediated by an evolutionarily conserved enzyme, arginyltransferase1 (Ate1). The downregulation of Ate1, as seen in many types of cancer, prominently increases cellular tolerance to a variety of stressing conditions. Conversely, in yeast and mammalian cells, Ate1 is elevated under acute oxidative stress conditions and this change appears to be essential for triggering cell death. However, studies of Ate1 were conventionally focused on its function in inducing protein degradation via the N-end rule pathway in the cytosol, leading to an incomplete understanding of the role of Ate1 in cell death. Our recent investigation shows that Ate1 dually exists in the cytosol and mitochondria, the latter of which has an established role in cell death initiation. Here, by using budding yeast as a model organism, we found that mitochondrial translocation of Ate1 is promoted by the presence of oxidative stressors and is essential for inducing cell death with characteristics of apoptosis. Also, we found that Ate1-induced cell death is dependent on the formation of the mitochondrial permeability pore and at least partly dependent on the action of mitochondria-contained factors including the apoptosis-inducing factor, but is not directly dependent on mitochondrial electron transport chain activity or its derived reactive oxygen species (ROS). Furthermore, our evidence suggests that, contrary to widespread assumptions, the cytosolic protein degradation pathways including ubiquitin-proteasome, autophagy, or endoplasmic reticulum (ER) stress response has little or negligible impacts on Ate1-induced cell death. We conclude that Ate1 controls the mitochondria-dependent cell death pathway.

## INTRODUCTION

The regulation of programmed cell death (PCD) in eukaryotes is essential for the organism to properly respond to stress conditions, to restrict the temporal impact of damages, to adapt to chronic stress, and to shape the tissues during development and regeneration. As such, the understanding of PCD events possesses the utmost significance for understanding eukaryotic life forms. In human, dysregulation of PCD is directly responsible for many types of diseases or abnormalities such as cancer, diabetes, autoimmunity, aging-related disorders, prenatal deformation, and embryonic lethality.

Studies from us and other groups showed that eukaryotic PCD is influenced by arginyltransferase (*ATE*) family proteins, which is known for being the only enzyme family mediating arginylation in eukaryotes. Arginylation is the posttranslational addition of an extra arginine to target proteins independently of the ribosome and mRNA[1]. This modification may take place on an eligible N-terminus or side chain of substrate proteins. By adding a charged and bulky chemical group, arginylation is expected to lead to significant changes of the properties of the substrate protein. When taking place at the N-terminus, arginylation often leads to hyper-ubiquitination and/or rapid degradation of the target protein by the proteasome or autophagy system via the N-end rule pathway[2, 3]. In all eukaryotes, the *ATE* genes are coded in the nuclear genome. While in plants the *ATE* family include two isoforms (Ate1 and Ate2) due to gene duplication, most other eukaryotes (such as fungi, animals, and humans) only carry one isoform, Ate1.

The protein level and/or the activity of Ate1 protein were found to increase upon the acute exposure to a broad range of stressors including oxidants, heavy metal, physical injury, and radiations[4-9]. Conversely, the decreased expression or inhibition of Ate1 often renders the cells insensitive to those stressing factors[9-11]. These data suggested that Ate1 may have a direct role in regulating cell death. Indeed, our past study showed that an ectopic overexpression of Ate1, even in the absence of exogenous stressors, is a potent inducer of cell death[9]. Consistently, multiple genetic studies uncovered many Ate1-associated phenotypes in embryo/tissue development and/or diseases that can be attributed to PCD regulation. For example, the knockout (KO) of Ate1 gene (*ATE1*) leads to complete embryonic lethality at the early-mid gestation stage [12]. Conditional deletion or dysregulation of *ATE1* also causes problems in cardiovascular development, neuronal development, and angiogenesis[13-16]. Furthermore, downregulation of Ate1 appears to contribute to the development and progression of cancer, which is often coupled with decreased PCD [11, 17].

In the attempts of addressing the role of Ate1/arginylation in PCD, two approaches have been used in past studies. In the first one, genetic manipulations or specific inhibitor were used to study the global consequences of interfering the level/activity of Ate1 and its functional partners. In most cases, the suppression of Ate1 or its functional partners (N-terminal amidohydrolase (NTAN) or ubiquitin-protein ligase E3 component n-recognin (UBR) drastically decreases cellular sensitivities to death-inducing stressors [10, 18-20], while direct upregulations of these corresponding elements often promote cell death[9]. As such, most of these data implied an overall pro-apoptotic role for Ate1/arginylation. In the second approach, researchers focused on identifying arginylation substrate proteins in the cytosol that are involved in PCD-related processes. These processes include protein degradation and/or quality-check machineries such as the ubiquitin-proteasome system (UPS), autophagy, and unfolded protein response (UPR) in the endoplasmic reticulum (ER)[21-23]. The identified Ate1 substrates include full-length proteins or proteolytic fragments generated by caspase or calpain. Many of the identified proteins/fragments appear to have known apoptosis-promoting roles (if present in the cytosol). Examples include BRAC1, RIPK1, and EPHA4. Since Ate1-mediated arginylation appears to accelerate their turnovers, this naturally led to the hypothesis of an anti-apoptotic role of Ate1/arginylation [24-27]. However, this speculation is at odds with the result from the first approach mentioned above. It is also at odds with other target-based studies including finding showing that arginylation of certain autophagy components (such as the cytosolic BiP) promotes autophagy and reduce PCD[22, 28]. These discrepancies indicated that the role of Ate1/arginylation in PCD is likely more complex than previously thought and that there is likely a major missing mechanistic link between the different approaches.

A new insight for the investigation of Ate1 was provided in our recent study showing a previously overlooked connection between Ate1 and mitochondria, the latter of which is a major regulator of PCD. In general, cell death can be triggered by the presence of stressors such as reactive oxygen species (ROS) or DNA damages, or by the signalling relayed by receptors for extracellular ligands (such as cytokines). These stimuli further activate intracellular signalling/actioning proteins such as the caspases to initiate the PCD process. A major regulator of PCD is mitochondria, which are the main generators of bioenergetics molecules (ATP) as well as endogenous ROS as a byproduct of electron transport chain (ETC) activity. Furthermore, a variety of stressing conditions or signalling events can cause mitochondrial membrane leakage, leading to the release of otherwise mitochondria-contained pro-death factors such as cytochrome *c* (Cyc), apoptosis-inducing factor (Aif), and endonuclease G (Nuc1). The release of these factors to the cytosol often triggers cell death events in caspase-dependent and independent manners[29, 30]. Our analysis revealed that the eukaryotic *ATE1* gene, which is hosted in the nuclear genome, was likely derived from mitochondrial gene transfer during early eukaryote evolution. We also found that, while the absolute majority of Ate1 resides in the cytosol and nucleus, a small fraction of this protein is located inside mitochondria[31]. Consistently, studies from us and others showed that Ate1 is essential for maintaining normal mitochondrial respiration and/or morphology in mammalian and yeast cells[31, 32]. Considering the known role of mitochondria as a central hub for initiating and coordinating PCD, we were inspired to question whether Ate1 regulates PCD through mitochondria-dependent pathways. However, past studies about Ate1 were nearly exclusively examining its actions in the cytosol or nucleus, lending little clue to address the role of Ate1 in relation to mitochondria.

To test how Ate1 regulates cell death, in this study we took advantage of the budding yeast (*Saccharomyces cerevisiae*) as a test model, for which well-established assays for PCD and Ate1/arginylation are readily available. In this organism, PCD is mediated by an ensemble of machineries that are conserved among other eukaryotic organisms. As in other eukaryotes, PCD in yeasts can be induced by stressors, signalling molecules, and aging process [33-35], Furthermore, the test of PCD in yeasts can be facilitated by an already demonstrated assay using over-expressed Ate1 with clear and robust readouts[9]. Here, by using *S. cerevisiae* as test model, we found that exposure to oxidative stressors lead to Ate1 translocation to mitochondria and cell death induction with characteristics consistent to apoptosis. Furthermore, Ate1-mediated PCD requires the opening of the mitochondrial permeability transition pore (mPTP) leading to mitochondrial outer membrane permeabilization (MOMP), and the presence of mitochondrial apoptogenic protein Aif1. However, mitochondrial Ate1 does not appear to directly induce ETC alterations or increase in ROS generation. Vice versa, ETC activity and ROS are not directly required for the execution of Ate1-indcued PCD. Finally, the execution of Ate1-mediated cell death is largely independent of most cytosolic PCD-relevant mechanisms including UPS, autophagy, and unfolded protein response (UPR). Therefore, our data suggests that Ate1 is an ancient and conserved regulator of the mitochondrial cell death pathway that was previously overlooked.

## RESULTS

### Ate1 is enriched in mitochondria upon stress treatments and induces cell death

In our previous study, we observed a trace amount of Ate1 protein located inside the mitochondria in both yeast and mammalian cells. As a first test to see how this localization is related to cell death, we examined the distribution of Ate1 in the presence or absence of oxidizing chemicals including hydrogen peroxide (H_2_O_2_) and sodium azide (NaN_3_), which are potent agents to induce PCD. To facilitate the tracing, the endogenous Ate1 in the yeast genome was fused with a GFP tag, which by itself does not appear to affect the function or location of Ate1 as shown in previous publications from us and others[9, 17, 36]. In absence of a stressor, the signal of Ate1 appears to be largely diffusive, as detected by microscopy. However, upon the exposure to oxidizing agents (H_2_O_2_ or NaN_3_), the Ate1-GFP signal rapidly enriches to puncta-like structures within 10 minutes (Fig 1A). These structures colocalize with the Mitrotracker-Red signal, a marker for mitochondria (Fig.1B), suggesting that Ate1 translocate to mitochondria in response to oxidative stress. To further quantitatively validate this change in Ate1 localization upon stress, we measured the Pearson colocalization coefficient of the Ate1-GFP and mitochondrial signals by microscopy. We found significant increased colocalization in stressor treated cells compared to the non-stressed cells (Fig 1C).

**Figure 1.**
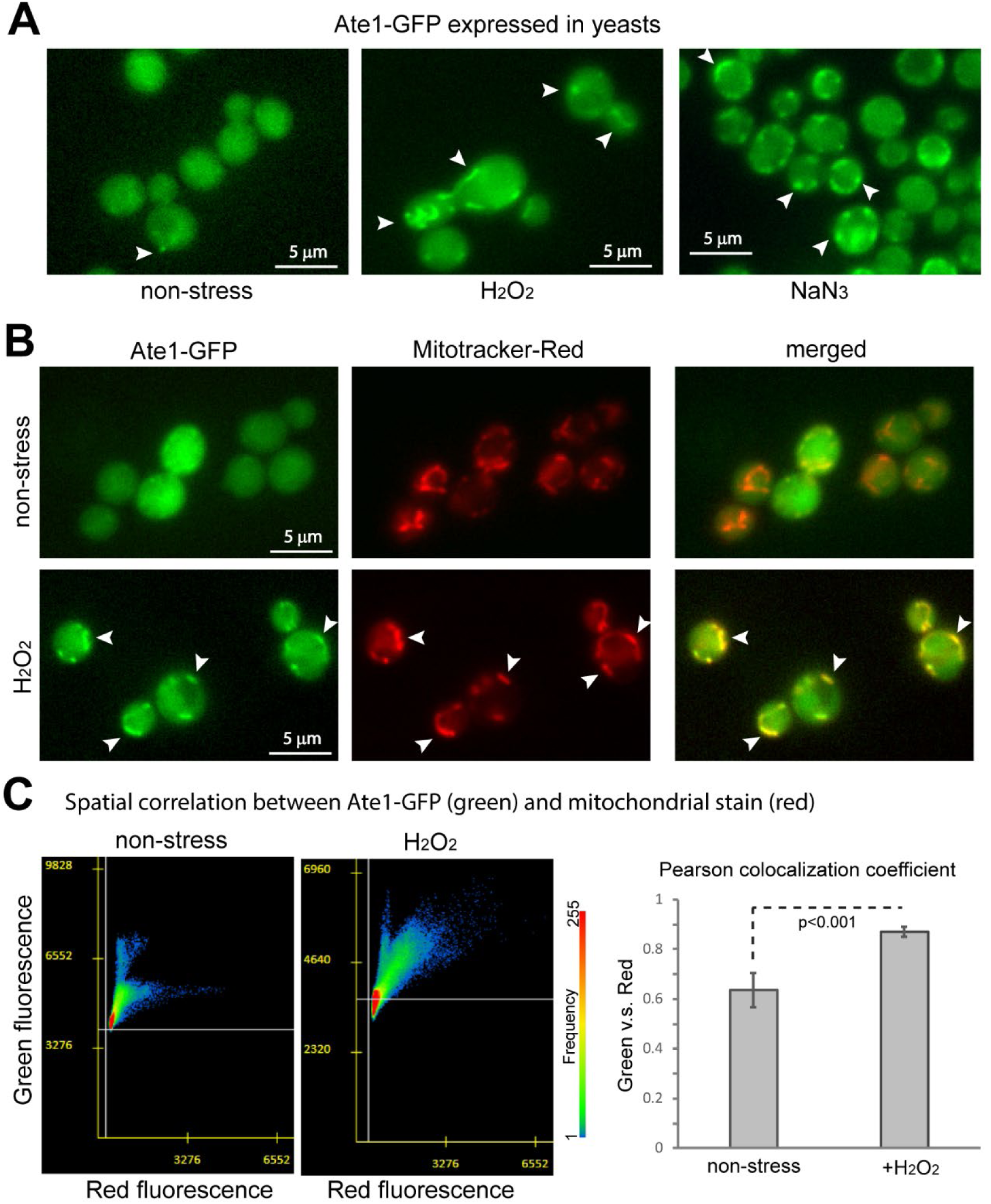
Visualization of Ate1 colocalization with mitochondria upon treatments of oxidative stressors. (A) Representative fluorescence microscopic images showing the signal of Ate1 tagged with GFP, which is driven by the endogenous *ATE1* promoter at the native chromosome locus. The yeast cells were either treated with 5mM H_2_O_2_ or 5% NaN_3_, or not treated with anything (nonstress). The observation started promptly within 5 minutes of treatments. The BY4741 strain yeast was used for tests in all figures unless otherwise indicated. (B) The above mentioned Ate1-GFP expressing yeast cells were treated with stressors H_2_O_2_ (5mM) and stained with mitochondria-specific dye Mitoracker-Red. Arrow point to several locations where obvious colocalization of the green and red fluorescent signals are observed. (C) To quantify the mitochondrial translocation of Ate1 under stressing conditions, Pearson colocalization coefficient analysis was used to measure the relative colocalization of GFP tagged Ate1 and mitochondria-specific dye (Mitotracker-red) in non-stressing and stressing conditions.

To corroborate the microscopy-based observations, we utilized an ectopic expression of a GFP-tagged recombinant Ate1 driven by a galactose-inducible promoter. The over-expressed Ate1 has been shown to increase cellular sensitivity to stressors and directly induce PCD [9, 37]. To avoid the interference of the endogenous Ate1, the recombinant protein was expressed in *ate1*Δ yeasts. To focus on the early event of PCD, the cells were analyzed at 6 hours post induction, before Ate1-mediated PCD induction[9]. We isolated mitochondria from yeast cells, incubated them with proteinase K to remove proteins attached to the mitochondrial outer membrane, and analyse Ate1 protein levels by immunoblotting. To test the impact of stress treatment in the mitochondrial import of Ate1, we compared yeast cells treated or not treated with the oxidative stressor NaN_3_. We detected Ate1-GFP in the mitochondrial fraction of both NaN_3_ treated and untreated samples. In agreement with our microscopy-based data, we observed that, while the stress treatment did not increase the overall level of recombinant Ate1 in the whole cell (Fig.2A), however, the mitochondria-specific fraction of Ate1 is significantly increased in the stressed cells (Fig.2B).

**Figure 2.**
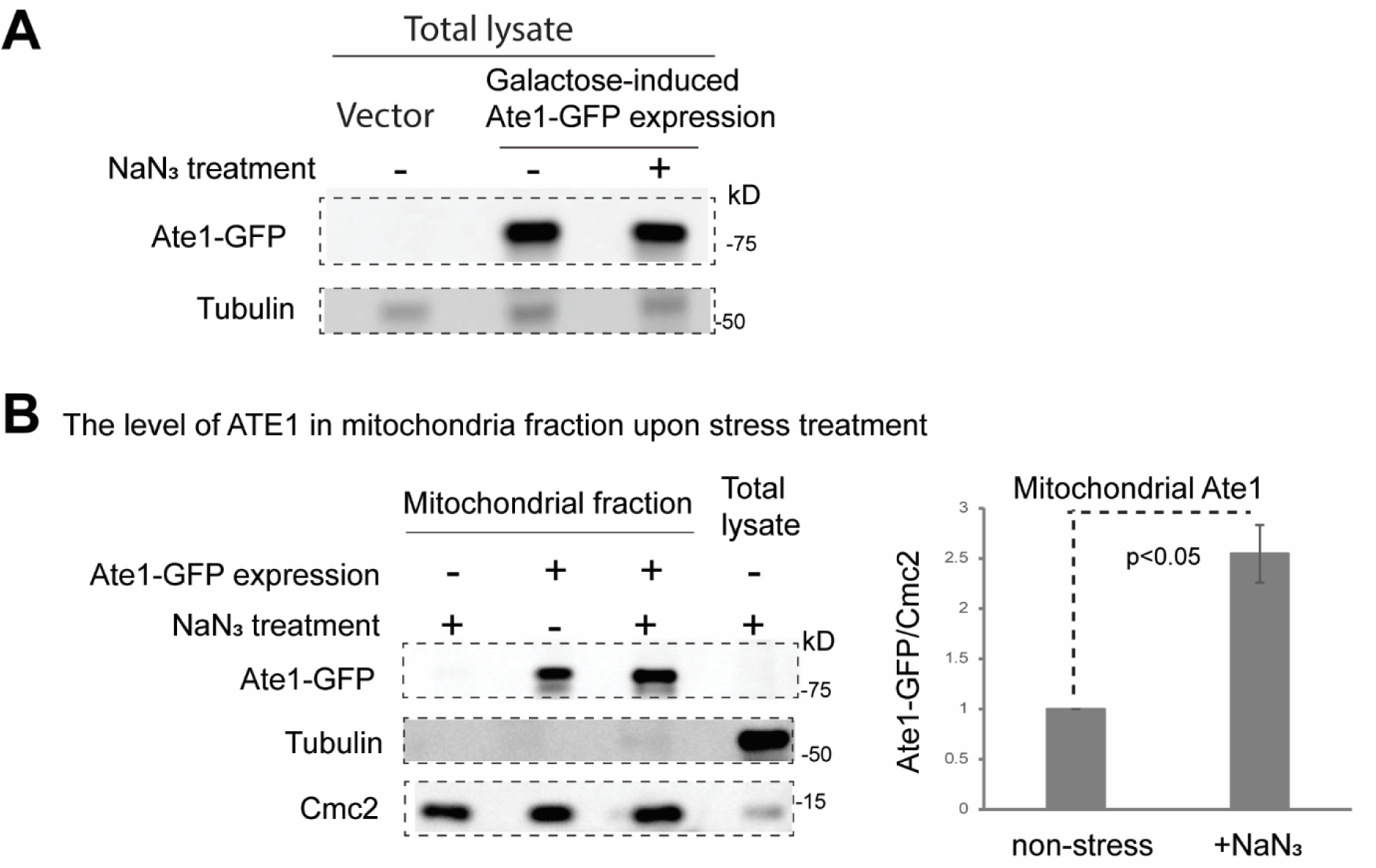
Oxidative stress induced by NaN3 increases mitochondrial Ate1 localization. (A) Representative Western blot showing total levels of Ate1-GFP in W303 wild-type strain treated either with or without 5%NaN_3_ for 30 minutes. Ate1-GFP expression was induced by the addition of 0.5% galactose and incubation for 3 hours at 30°C. Empty vector serves as a negative control. Alpha tubulin is used for protein loading control. (B) Left panel; representative Western blot showing Ate1-GFP levels in purified mitochondrial fractions, which was treated with proteinase K (5 μg/ml) to remove proteins that are not protected by the mitochondrial membrane. Alpha-tubulin is used as a marker for cytosolic protein contamination, while mitochondrial intermembrane space protein Cmc2 serves as a marker for mitochondrial proteins. Right panel: showing the quantification of mitochondrial Ate1-GFP levels normalized to mitochondrial protein Cmc2. Error bar denotes SD (N=4), p values were calculated by two tailed student t-test.

To further test if the mitochondrial localization of Ate1 is required for the induction of cell death, we over-expressed Ate1 for an extended duration, a condition that can cause PCD in the absence of exogenous stressor[9]. As a control, to prevent Ate1 mitochondria localization, we fused the Ate1-GFP with an additional tag derived from the Ras2 protein, which is expected to be conjugated to lipid molecule on the plasma membrane[38]. As validated by microscopy, the expressed Ate1-GFP-Ras2 localizes to the periphery of the cells even under stress, and does not form any puncta-like structures (characteristic of mitochondria) unlike to the Ate1-GFP (Fig 3A). To test if the Ras2 tag negatively affects the activity of Ate1, we used an arginylation reporter, DD-β15, which is derived from the N-terminal sequence of a known arginylation substrate β-actin[9, 39]. It is fused to a fluorescence protein mCherryFP, which is expected to be diffusively distributed in the cytosol and nucleus (Fig.3B). By using the co-expression of the reporter with the recombinant Ate1, we found that, while a lower expression level of Ate1 is seen with the Ras2-tagging, the arginylation level of the reporter remains similar (Fig.3C). This indicates that the Ras2 tagging did not decrease the arginylation activity of the recombinant Ate1 inside the cytosol. However, in contrast to the original Ate1-GFP, the overexpression of Ate1-GFP-Ras2 leads to a higher viability of yeast cells (Fig 3D), suggesting that blocking the mitochondrial localization of Ate1 minimizes its effects in inducing cell death. To test this possibility from a reverse angle, we tagged the Ate1 with a mitochondrial targeting signal and found it exhibited a more potent effect for inducing cell death compared to the original Ate1-GFP (Fig. 3 E & F). Therefore, our results together showed that the mitochondrial localization of Ate1 is required for Ate1-induced cell death.

**Figure 3.**
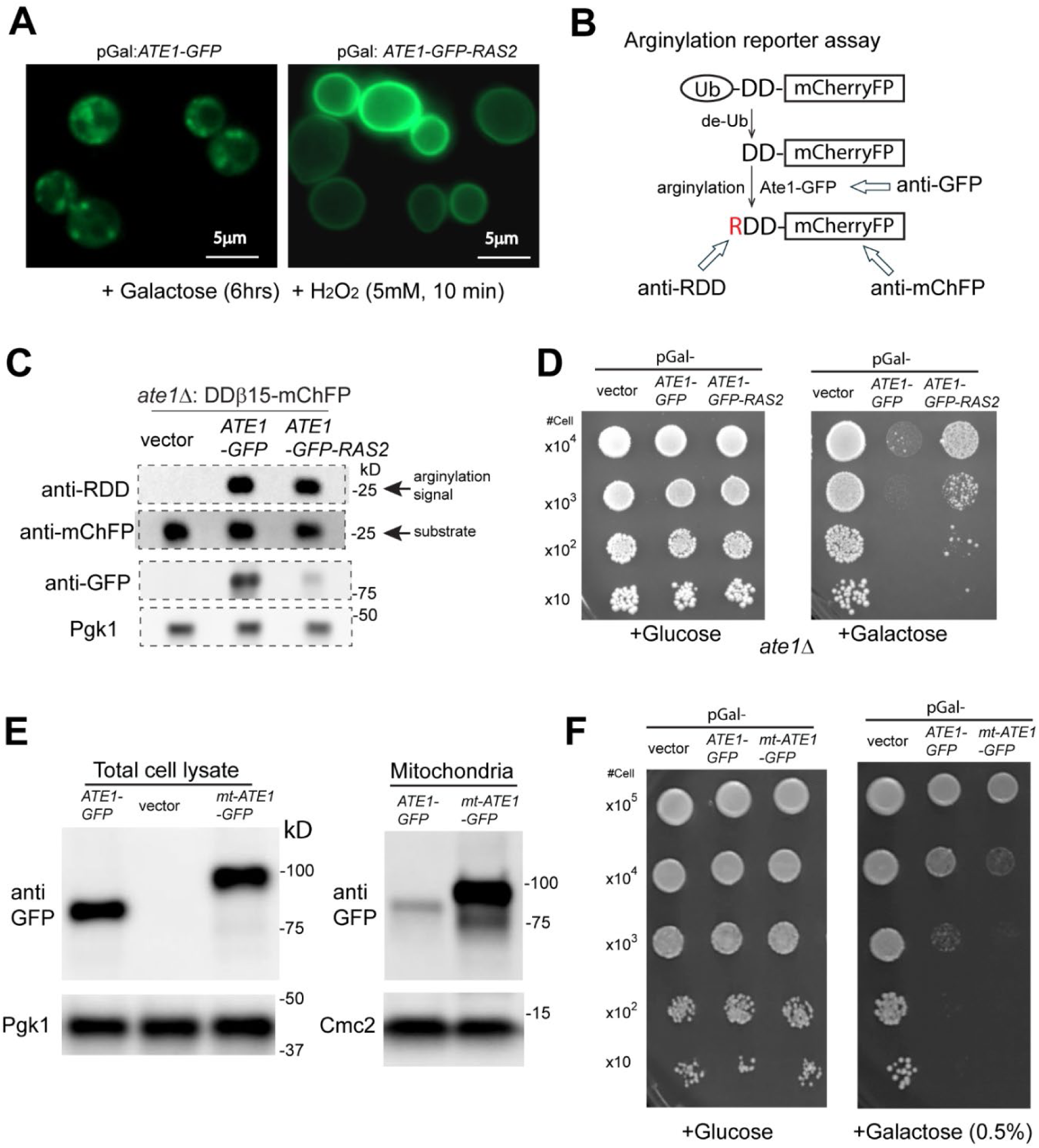
The mitochondrial translocation of Ate1 is essential for the cell death induced by Ate1 over-expression. (A) Yeast cells containing Ate1-GFP and Ate1-GFP-Ras2 that are driven by galactose-inducible promoters (pGal) were allowed to express for 6 hours and then briefly treated with oxidative stressor H_2_O_2_ for 10 minutes before the microscopic images were taken for the green fluorescence channel. The regular Ate1-GFP forms puncta-like structures similar as shown in Fig.1, while Ate1-GFP-Ras2 remains localized to the periphery of the cells. (B) Scheme illustrating the principle of the reporter for the arginylation activity inside yeast cells. The N-terminal ubiquitin domain of the reporter protein DD-β15-mCherryFP will be promptly removed by endogenous de-ubiquitination (de-Ub) enzymes in the cell, exposing the penultimate peptide DD-β15, which is derived from the N-terminus of mouse β-actin and is known to be arginylated *in vivo*[9, 39]. The arginylated N-terminus can be recognized with a specific antibody anti-RDD[9]. Antibodies for mCherryFP (mChFP) and GFP can be used to probe the levels of the reporter protein and the GFP-fused ATE1, respectively. (C) To test the arginylation activity of different forms of Ate1 (Ate1-GFP or Ate1-GFP-Ras2), they were expressed in *ate1*Δ yeast (to avoid the interference of endogenous ATE1), which was also simultaneously expressing the reporter protein DD-β15-mCherryFP. The arginylation level of the reporter protein was measured as described in (B). Pgk1 serves as loading controls for the yeast proteins. (D) Growth test of *ate1*Δ yeast cells carrying either the empty expression vector or pGAL1:Ate1-GFP and pGAL1:Ate1-GFP-Ras2 was conducted by a serial dilution growth assay on either plate containing glucose or galactose, where the expression of Ate1 is not induced or induced, respectively. (E) Representative Western blots showing the expression levels and distributions of different Ate1 constructs (Ate1-GFP and the mitochondrial matrix-targeting Mt-Ate1-GFP) in total cell lysate (left panel) and in purified mitochondrial fractions (right panel). Vector alone served as a negative control. The expression of both the Ate1 construts was achieved by adding 0.5% galactose and incubated for 3hrs at 30_o_C. The expressions and subcellular distributions Ate1-GFP and Mt-Ate1-GFP were probed by anti-GFP. The level of Pgk11 and Cmc2 were used as loading controls for total proteins or mitochondrial proteins, respectively. (F) Growth test of *ate1*Δ yeast cells carrying either the empty expression vector, pGAL1:Ate1-GFP, or the mitochondria matrix localized Ate1 (pGAL1: Mt-Ate1-GFP) by a serial dilution growth assay on either plate containing galactose or glucose for the induced expression (or not) of Ate1. Note that the concentration of galactose (0.5%) is lower than elsewhere to allow the display of the difference between the Ate1-GFP and mt-Ate1-GFP. Plates were incubated at 30°C and images were taken after 3 days.

### Ate1-induced cell death is dependent on mitochondrial permeability transition pore with characteristics of apoptosis

Similar to what is seen in metazoan, yeast mitochondria play a central role in several PCD scenarios such as apoptosis, necrosis, and pyroptosis. To distinguish these pathways, we used the TUNEL assay to examine yeasts with over-expressed Ate1 (fused with a non-fluorescence tag 6xHis). We found the Ate1-induced cell death is accompanied with positive signs of large-scale DNA cleavages as shown in the TUNEL assay (Fig.4A), which is suggestive of apoptosis[40].

**Figure 4.**
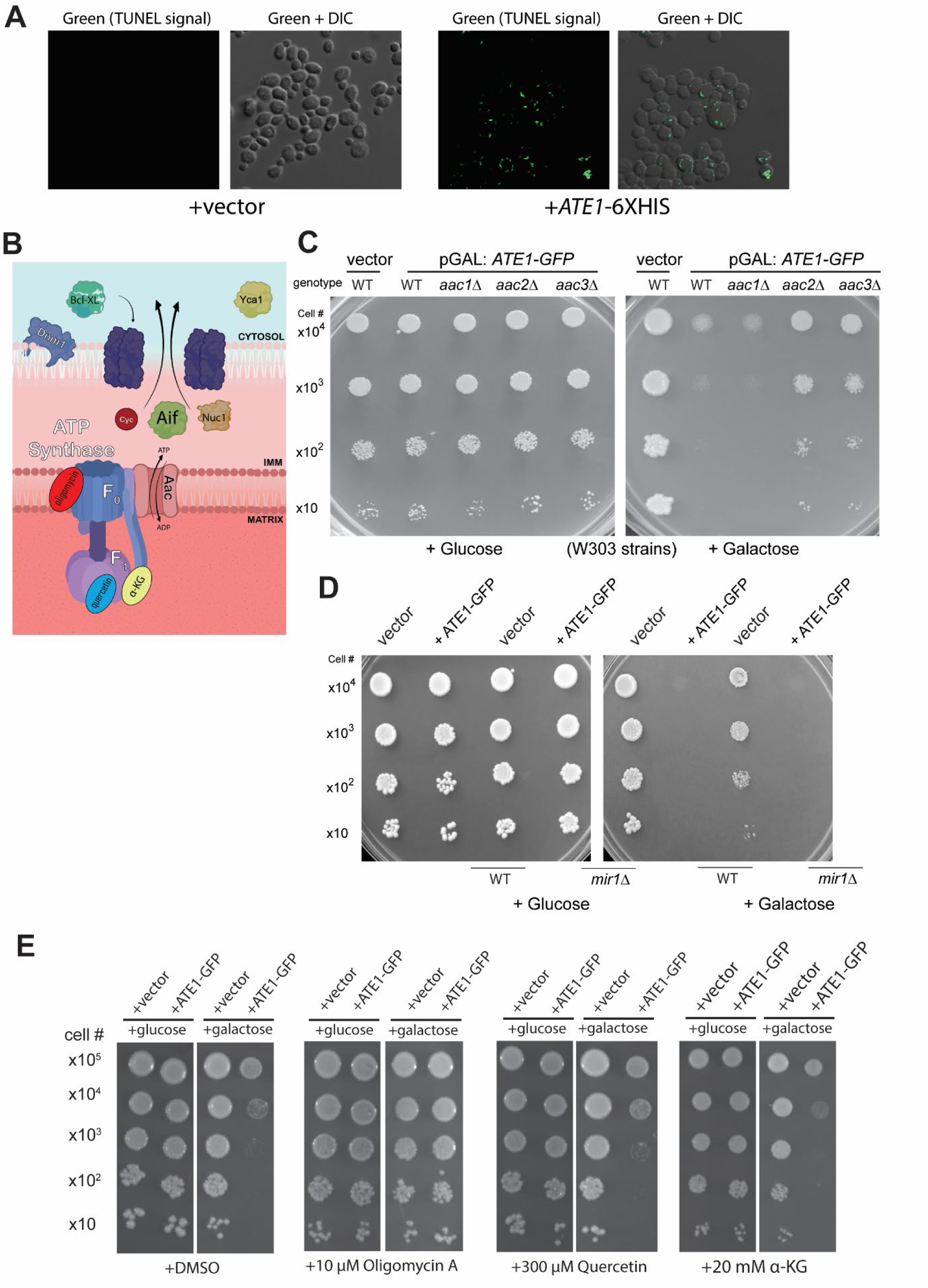
Ate1-overexpression triggers apoptosis with a dependency on mitochondrial permeability transition pore (mPTP) A) Representative microscopic images displaying the presence of DNA fragmentations probed by the TUNEL assay on *ate1*Δ yeasts that were induced by galactose for the expression of Ate1-6xHIS (or the empty vector). The panels on the left display only the TUNEL signal (green fluorescence), while the panels on the right are an overlay of TUNEL and Differential Interference Contrast (DIC) microscopic images. B) A diagram showing some of the key components in yeasts that affect mPTP and apoptosis. Note that the Bcl-XL is not an endogenous protein of yeast, but it can interact with the mitochondrial outer membrane permeabilization (MOMP) event[50]. C) Growth of yeast cells (WT, *aac1*Δ, *aac2*Δ, *aac3*Δ, all on W303 strains) carrying either the empty expression vector (vector) or pGAL1:Ate1-GFP (+Ate1-GFP) was measured by a serial dilution growth assay on either plate containing 2% glucose or 2% galactose, where the expression of Ate1 is not induced or induced, respectively. Plates were incubated at 30°C and images were taken after 2-3 days. D) Similar to (B), except that WT and *mir1*Δ yeasts were used. E) Serial dilution growth assay to assess changes in growth of WT yeast induced with pGAL: Ate1-GFP in the context of various ATP synthase inhibitors. Yeasts were grown in raffinose-containing liquid media before being washed, serially diluted in H_2_0, and plated to either glucose or galactose-containing plates with the designated concentrations of inhibitors. Plates were allowed to grow three days before images were taken. DMSO (at a final concentration < .004%) was used as vehicle control for Oligomycin A and Quercetin.

In most eukaryotes, the mitochondrial apoptotic pathway is driven by the formation of the mitochondrial permeability transition pore (mPTP), a large, unselective channel present within the inner membrane, which leads to the swelling of mitochondria and rupture of the mitochondrial outer membrane. It is important to point out that the molecular composition of the mPTP has shown a degree of flexibility. However, some of the most conserved core elements include the F_o_ module of the F_o_F_1_-ATPase and isoforms of the adenine nucleotide transporter (ANT) (Fig. 4B)[29, 41-43]. To determine the involvement of these components in Ate1-induced apoptosis, we overexpressed Ate1 in yeast strains with deletions of the corresponding genes. We found that the deletions of two ANT isoforms (*AAC2* and *3*) were able to generate significant rescue effects on the Ate1-induced lethality (Fig. 4C). In addition to mPTP formation during apoptotic events, ANT normally facilitates the exchange of nucleotides (ATP/ADP) coupled with ATP synthesis. To exclude the possibility the later role is involved in Ate1-induced cell death, we examined the knockouts of *MIR1*, which is involved in mitochondrial phosphate transport (important for ATP synthesis) but not directly involved in mPTP formation[29]. We found that this gene knockout does not generate rescue effects on cell death induced by Ate1 overexpression (Fig. 4D). To corroborate with the above genetics-based tests, we used specific inhibitors to block the functions of F_o_F_1_-ATPase, another important component of mPTP. We found that, the application of oligomycin, which specifically blocks the function of the F_o_ subunit of F_o_F_1_-ATPase[44], potently supresses the lethal effects of Ate1-overexpression (Fig. 4E). However, in addition to block the formation of the mPTP, oligomycin also would compromise ATP synthesis. To exclude the latter possibility, we applied two different reagents, quercetin and α-ketoglutarate, which inhibit the F1 compartment of ATPase to impair ATP synthesis but not the mPTP formation[45, 46]. We found these treatments has little effects on the Ate1-induced cell death (Fig. 4E). These data suggests that Ate1 specifically regulates the mitochondrial mPTP formation for the induction of apoptosis.

The formation of the mPTP is usually further connected with the mitochondrial outer membrane permeabilization (MOMP) events, which leads to the release of apoptogenic proteins such as cytochrome *c* (Cyc), apoptosis-inducing factor (Aif1), and Endonuclease G (Nuc1) from the mitochondrial intermembrane space to the cytosol/nucleus, where they contribute to the apoptosis execution. To test the relevance of MOMP in Ate1-induced cell death, we utilized exogenous expression of mammalian Bcl-xL, a Bcl-2 family protein that is known to block the outer channel of MOMP formed by both Bax-dependent[47] and independent mechanisms[48, 49]. Although yeast lacks Bcl-2 family proteins, it contains the other mPTP and MOMP components that were shown to react to mammalian Bcl-2 in a similar fashion [50]. To test the possibility that Ate1 induced MOMP can be prevented by Bcl-xL, we co-expressed this protein in yeast cells with the over-expressed Ate1. We found that Bcl-xL expression indeed has measurable effects in antagonising Ate1-induced lethality, similar to its effect found in mammalian Bax-induced yeast cell death (Fig. 5A). To directly test if over-expressed Ate1 causes MOMP, we examined the distribution of cytochrome *c* between mitochondria and the cytosol. Indeed, we found a higher portion of cytochrome *c* present in the cytosol *v.s.* mitochondria in yeast cells with over-expressed Ate1 (Fig. 5B), indicating a leakage of mitochondrial contents. Furthermore, as another validation of the relevance of MOMP events in Ate1-induced cell death, and to delineate the role of different mitochondria-containing apoptogenic factors, we used yeast with deletions of those corresponding genes including *AIF1* and *NUC1*. Considering yeast contains two *CYC* genes, we used the deletion of *YCA1*, which codes for the metacaspase that is supposed to respond to Cyc. We also used the deletion of *CYC3*, which codes for the lysase for the heme loading for the functional cytochrome c. We found that, while we observed no significant rescue by the deletions of *YCA1*(Fig.5D), *CYC3* (Fig.5E), or *NUC1* (Fig.5D), *AIF1*-knockout generated measurable rescue effects on Ate1-induced cell death (Fig.5 C&D), which supports the role of Ate1 in MOMP as a part of the mechanism of Ate1-mediated PCD. In comparison, the deletion of Dynamin-related GTPase (*DMN1*), which typically mediates mitochondria fission and receives signal from the cytosol and the cell membrane to trigger apoptosis, shows no effects (Fig.5 D).

**Figure 5.**
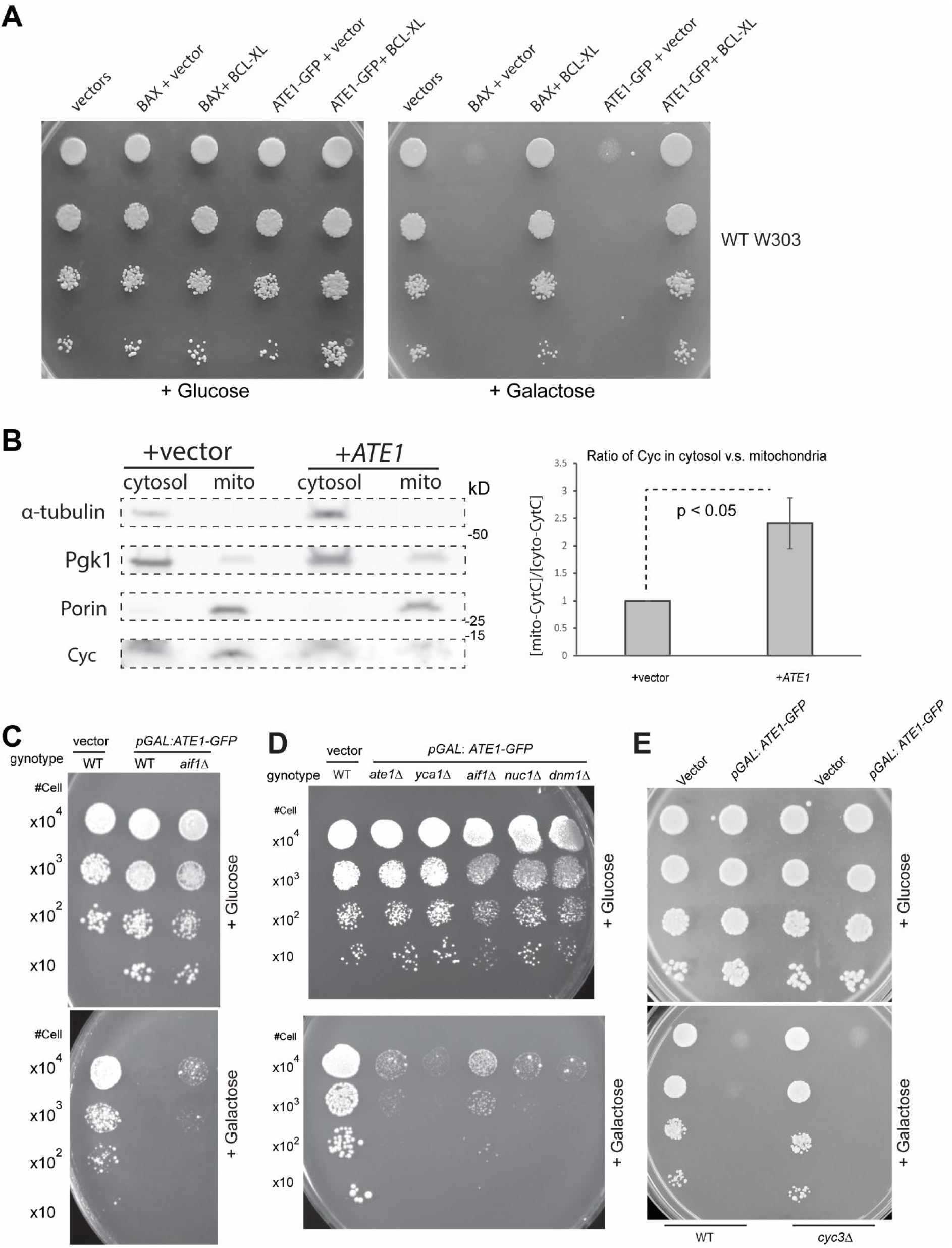
Ate1-induced cell death involves mitochondrial outer membrane permeabilization. A) Growth of yeast cells (W303 strain, WT) carrying either the empty expression vectors (pYES2-URA3 and pBF339-TRP1 vectors), or the galactose-inducible mouse BAX (pBM272-pGAL-*BAX*-URA3) and yeast Ate1 (pYES2-pGAL-*ATE1-GFP*-Ura3) in the presence of constitutively expressing Bcl-xL (pBF339-ADH-*BCLxL*-TRP1) or the vector (pBF339-TRP1) was measured by a serial dilution growth assay on either plate containing 2% glucose or 2% galactose, where the expression of Ate1 or Bax is not induced or induced, respectively. Plates were incubated at 30°C and images were taken after 3 days. B) Representative Western blots displaying changes in the distribution of cytochrome c (Cyc) between mitochondria and cytosol in yeast cell where Ate1-6xHis was induced for expression with 2% galactose in Ura-minus liquid media for 6 hours at 30°C. The separated cytosolic fraction and mitochondrial fraction were compensated by buffer to be equal volume before analysis. Alpha-tubulin and Porin (VDAC) were used as markers to display the purity of cytosolic and mitochondrial fractions, respectively. Pgk1 was used as a loading control. The quantification of the blot was based on n = 6; error bars represent S.E.M. C) Growth of yeast cells (WT or *aif1*Δ) carrying either the empty expression pYES2 vector (vector) or pYES-pGAL1:*ATE1-GFP* (+Ate1-GFP) was measured by a serial dilution growth assay on either plate containing 2% glucose or 2% galactose, where the expression of Ate1 is not induced or induced, respectively. Plates were incubated at 30°C and images were taken after 3-5 days. D) Similar to (C), except that WT, *aif1*Δ, *ate1*Δ, *yca1*Δ, *nuc1*Δ, and *dnm11*Δ) yeasts were used. E) Similar to (C), except that WT and *cyc3*Δ yeasts were used.

Therefore, our data together suggest that the elevation of Ate1 induces apoptosis by an intrinsic, mitochondria-dependent pathway.

### Mitochondrial electron transport chain (ETC) and mitochondria-generated reactive oxygen species (ROS) are not directly required for the execution of Ate1-induced PCD

Mitochondrial electron transport chain (ETC) activity and mitochondria**-**generated reactive oxygen species (ROS) have been shown to play important role in yeast apoptosis. Recent studies showed that an Ate1-deletion in either yeast or mammalian cells lead to a decrease in aerobic respiration[31, 51]. This prompted us to test if an elevation of Ate1 induces a hyperactivation of mitochondrial respiration and/or ROS generation, which then can lead to apoptosis by other mechanisms including the activation of caspases.

To test if the ETC activity is required for Ate1-induced cell death, we used a respiratory deficient yeast strain due to the deletion of the RIP1 gene, which encodes for a catalytic subunit of complex III. While the deletion of RIP1 indeed affected mitochondrial membrane potential, as confirmed by two different membrane potential-dependent dyes (mitotracker-red and rhodamine 123), it did not rescue Ate1-induced cell death (Fig.6A &B). To validate this result, we examined the knockout of *NDI1*, which encodes for the mitochondrial internal NADH dehydrogenase, and whose deletion also significantly compromise ETC activity (Fig.6B). We found that, similar to the case of *RIP1*-deletion, the knockout of *NDI1* does not show any protection against Ate1-induced cell death. These results are consistent with the previously mentioned data regarding ATPase inhibition by α-ketoglutarate or quercetin (Fig.4E), as well as the lack of effects by the deletion of *CYC3*(Fig.5E), which is also essential for ETC function. Taken together, these data demonstrate that the execution of Ate1-mediated cell death does not directly requires the ETC activity.

**Figure 6.**
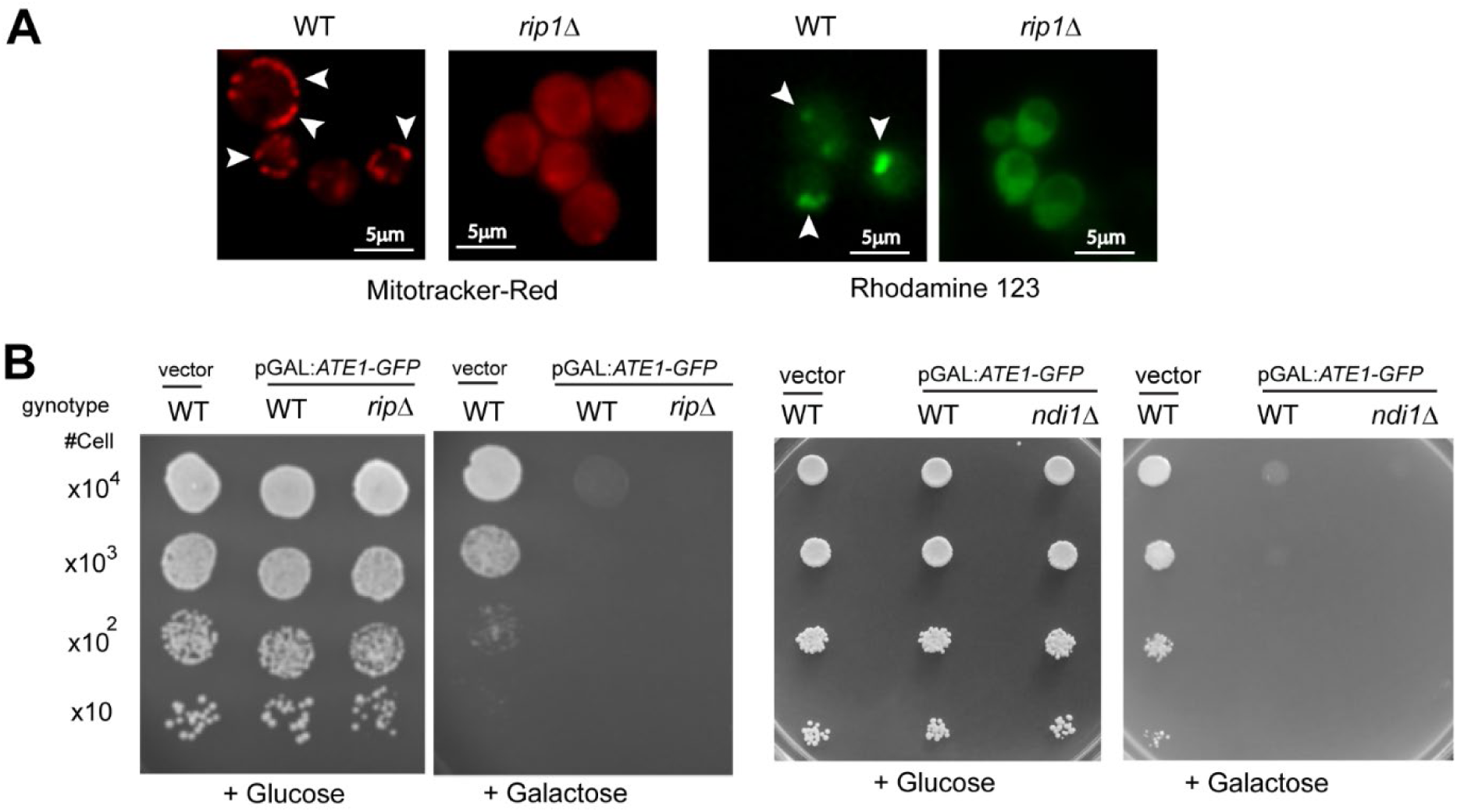
The mitochondrial respiratory activity is not directly required during Ate1-driven cell death. A) The membrane potentials in yeast cells, WT or *rip1*Δ, were probed by staining dyes Mitotracker-red (red) or rhodamine 123 (green). B) Growth of yeast cells (WT, *rip1*Δ or *ndi1*Δ) carrying either the empty expression vector pYES2 (vector) or pYES2-pGAL1:*ATE1-GFP* (+Ate1-GFP) was measured by a serial dilution growth assay on either plate containing 2% glucose or 2% galactose, where the expression of Ate1 is not induced or induced, respectively. Plates were incubated at 30°C and images were taken after 3-5 days.

To test if the elevation of Ate1 would trigger the generation of mitochondrial ROS, we measured the level of mitochondrial ROS in yeast over-expressing Ate1 in different timepoints (up to 12 hours, where the cell is expected to enter the apoptotic program[9]). However, we did not observe any significant increase in mitochondrial ROS levels throughout this time course (Fig.7A). To test this from a different angle, we co-expressed ROS neutralising enzymes superoxide dismutase (Sod1 or Sod2, which is located in cytosol or mitochondria, respectively) with Ate1. We found they generated no protecting effects in Ate1-induced cell death (Fig. 7B). Consistently, as mentioned earlier, the treatment of yeasts with quercetin, which is an inhibitor of mitochondrial ATP synthesis and also an anti-oxidant known to quench ROS in both mitochondria and cytosol, generates little effects on the Ate1-induced cell death. Therefore, while we previously showed that the presence of oxidative stress would upregulate Ate1 expression and would induce its translocation to mitochondria, the elevated Ate1 does not directly induced ROS production. Also, after Ate1 level is elevated, it does not require ROS to further carry on the cell death program.

**Figure 7.**
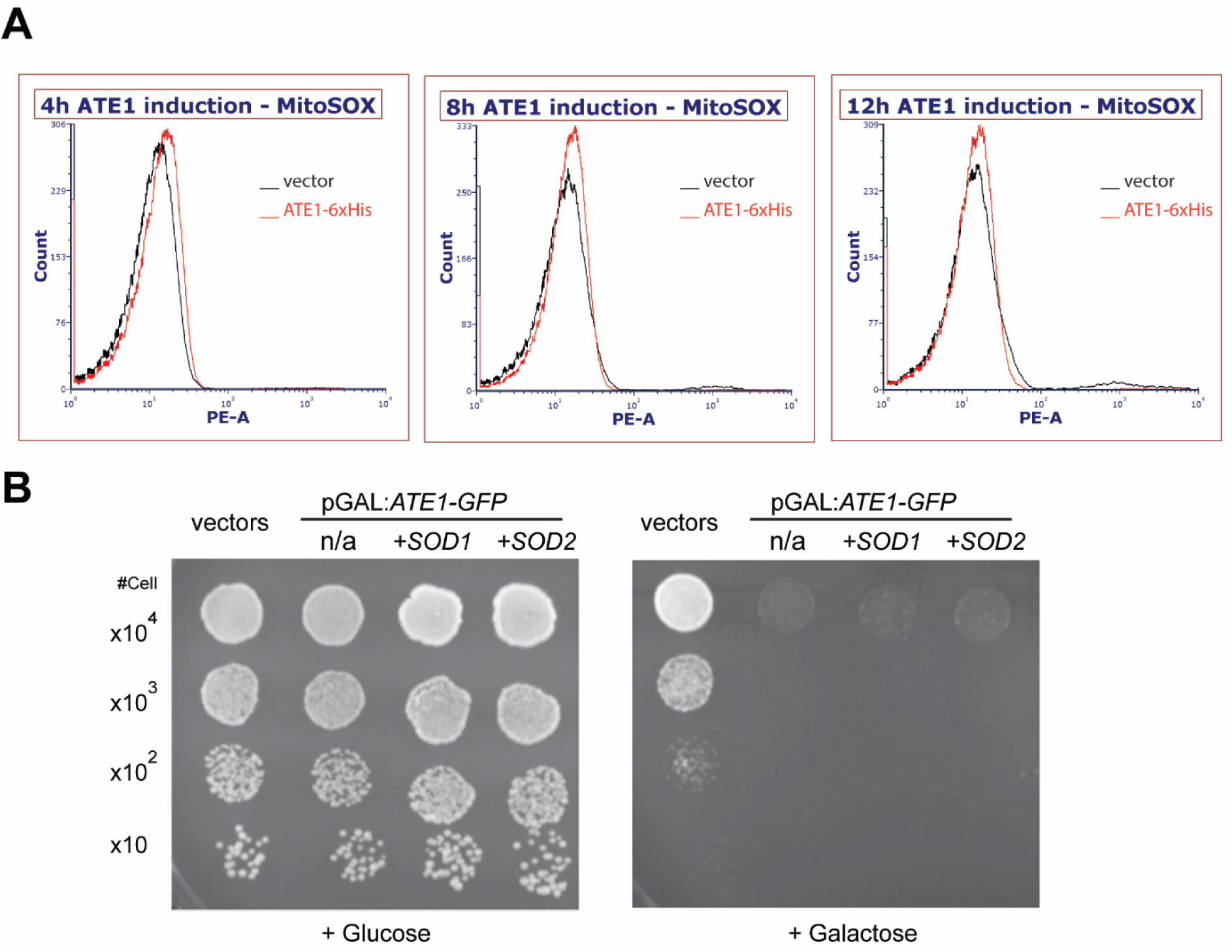
Mitochondrial ROS is not directly required during the Ate1-driven cell death. A) Representative flow cytometry plots depicting mitochondrial ROS levels at indicated timepoints after galactose induction. The yeast cells contain either the empty vector control pYES2 (black) or the pYES2-*ATE1*-6xHis for the overexpression of the 6xHis tagged Ate1 (red). MitoSOX Red was used to stain for mitochondrial ROS, and samples were excited with a 488 nm blue laser and emission was measured using a 585 nm filter with a 42 nm bandpass. No obvious differences in mitochondrial ROS were noted between Ate1 and control samples, in any of the time points. B) Growth of yeast cells (*ate1*Δ) carrying either the empty expression vectors pYES2-URA + pGPD2-Leu2 (vectors), pYES2-pGAL1:*ATE1-GFP* + pGPD2-Leu2 (n/a), or pYES2-pGAL1:*ATE1-GFP* in the presence of constitutively expression vectors of pGPD2-*SOD1*-Leu2 or pGPD2-*SOD2*-Leu2 (+ *SOD1*, or +*SOD2*). The growth rate was measured by a serial dilution growth assay on Ura-minus, Leu-minus SD plates containing 2% glucose or 2% galactose, where the expression of Ate1 is not induced or induced, respectively. Plates were incubated at 30°C and images were taken after 3 days.

### Cytosolic pathways including UPS, autophagy, and UPR are not required for the execution of Ate1-induced cell death

Ate1 was traditionally considered as a cytosolic protein and its functions were extensively characterized in the context of promoting protein degradations in relation to UPS, autophagy and UPR. As such, past studies relevant to Ate1 in cell death were mainly designed to test the effects of Ate1 on the cytosolic proteome. However, our above findings for the role of Ate1 in mitochondria-dependent pathway compelled us to re-examine these cytosolic pathways in the context of Ate1 and cell death.

Ate1-mediated arginylation was long considered a flagging signal for the recognition of UBR family E3 Ubiquitin (Ub) ligases. As such, an elevation of Ate1 would be expected to lead to an excessive ubiquitination, which in turn could trigger apoptosis by two scenarios. First, the presence of excessive ubiquitinated proteins may clog proteasome and/or autophagy systems, which can then cause apoptosis. Second, an over-activation of arginylation may lead to inadvertent depletion of proteins that are otherwise essential for maintaining cellular viability. To test the first possibility, we examined the global ubiquitination levels because an elevation of ubiquitin ladder/smear would be expected if the degradation machinery is clogged. However, we found that the over-expressing Ate1 did not lead to a significant increase of ubiquitin smear (Fig.8A). Consistently, when we examined the level of cytosolic heat-shock protein 70 (Hsp70), a marker for cytosolic unfolded protein stress, we found it was not increased, but moderately decreased when Ate1 is over-expressed (Fig.8B). Therefore, while our results did not rule out a role of Ate1 in ubiquitination, they clearly show that Ate1-overexpression did not lead to the clogging of UPS or other protein degradation pathways. In other words, the cause of Ate1-induced cell death is unlikely caused by the clogging of UPS. To test the second scenario, we used yeasts with the deletion of UMP1, which is expected to compromise proteasome activity and reduce protein turnover. However, this deletion does not provide any protective effects against Ate1-induced cell death (Fig.8C). As a further validation, we used yeast with the deletion of the only UBR enzyme (Ubr1) in yeast, which is expected to block the ubiquitination of arginylated proteins, and again we saw no rescue effects by this knockout (Fig.8C). Therefore, the cell death mediated by elevated Ate1 is not very likely to be mediated by an excessive degradation of cytosolic proteins.

**Figure 8.**
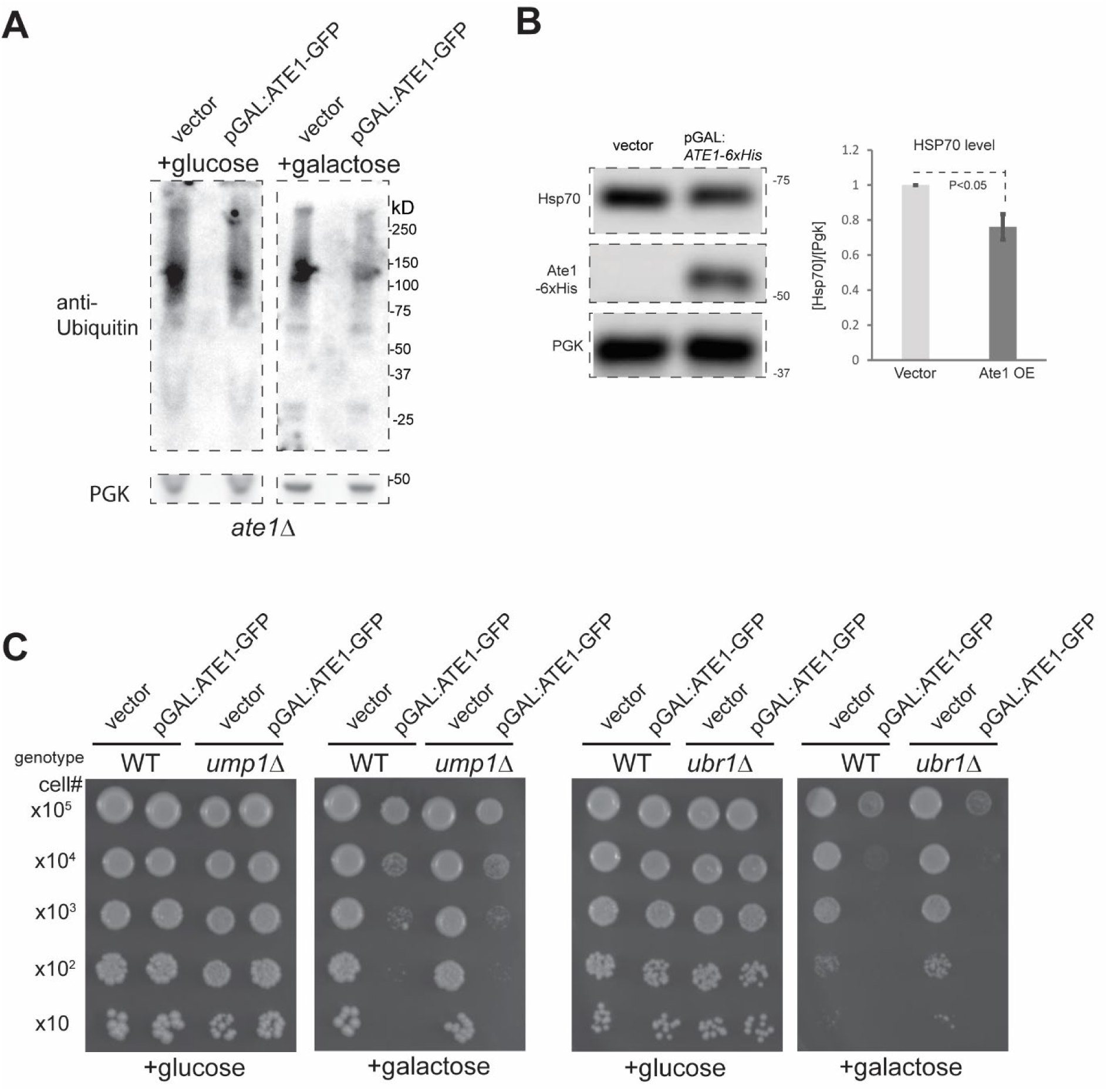
Ate1-overexpression does not lead to elevation of global ubiquitination and Ate1-induced cell death is not directly dependent on the functions of the ubiquitin-proteasome system. A) Representative Western blots depicting global ubiquitination levels in *ate1*Δ yeast with either pYES2-pGAL:*ATE1-GFP-*URA3 (pGAL:*ATE1-GFP*) or empty vector, which were induced for 6h with 2%galactose in liquid media. Pgk1 was used as a loading control. Images are from same gel. B) Representative Western blot showing the expression levels of cytosolic stress response protein HSP70 in the yeast cells carrying the pYES2-pGAL:*ATE1-6xHis-*URA3 expression vector or the empty control vector, which were induced with 2% galactose for 6h in liquid media. The level of Ate1 was probed with antibody against 6xHis tag. Right side panel shows the densitometric analysis of the cytosolic Hsp70 levels as expressed in fold difference after normalization to the internal protein loading control Pgk1. Error bar denotes SEM; N=6. C) Growth assay of WT, *ump1*Δ or *ubr1*Δ yeast carrying either pYES2-pGAL:*ATE1-GFP-*URA3 or empty vector. The growth was measured by a serial dilution growth assay on Ura-minus SD plates containing 2% glucose or 2% galactose, where the expression of Ate1 is non-induced or induced, respectively. Plates were incubated at 30°C and images were taken after 3 days.

Ate1/arginylation has been shown to interact with UPR and autophagy, both of which are known to regulate PCD. For example, an elevation and prolonging of UPR signalling is also known to initiate apoptosis and other PCD processes. The role of autophagy in PCD is less straight forward. While a normal level of autophagy is in general protective, a disruption or an over-activation of autophagy can lead to cell death. To test these possibilities, we examined the UPR stress by probing the level of GRP78, a marker of UPR signalling, and found that the over-expression of Ate1 leads to a significant reduction of UPR, suggesting that this is unlikely to be the direct cause of cell death in this scenario (Fig.9A). Furthermore, to probe the involvement of the autophagy, we used Atg8-GFP, whose cleavage is an established marker for autophagy activity. We found that Ate1-overexpression lead to negligible changes on autophagy activity, suggesting that this process is not directly involved (Fig.9B). Therefore, neither UPR or autophagy is likely to be the direct drivers of cell death induced by Ate1 elevation.

**Figure 9.**
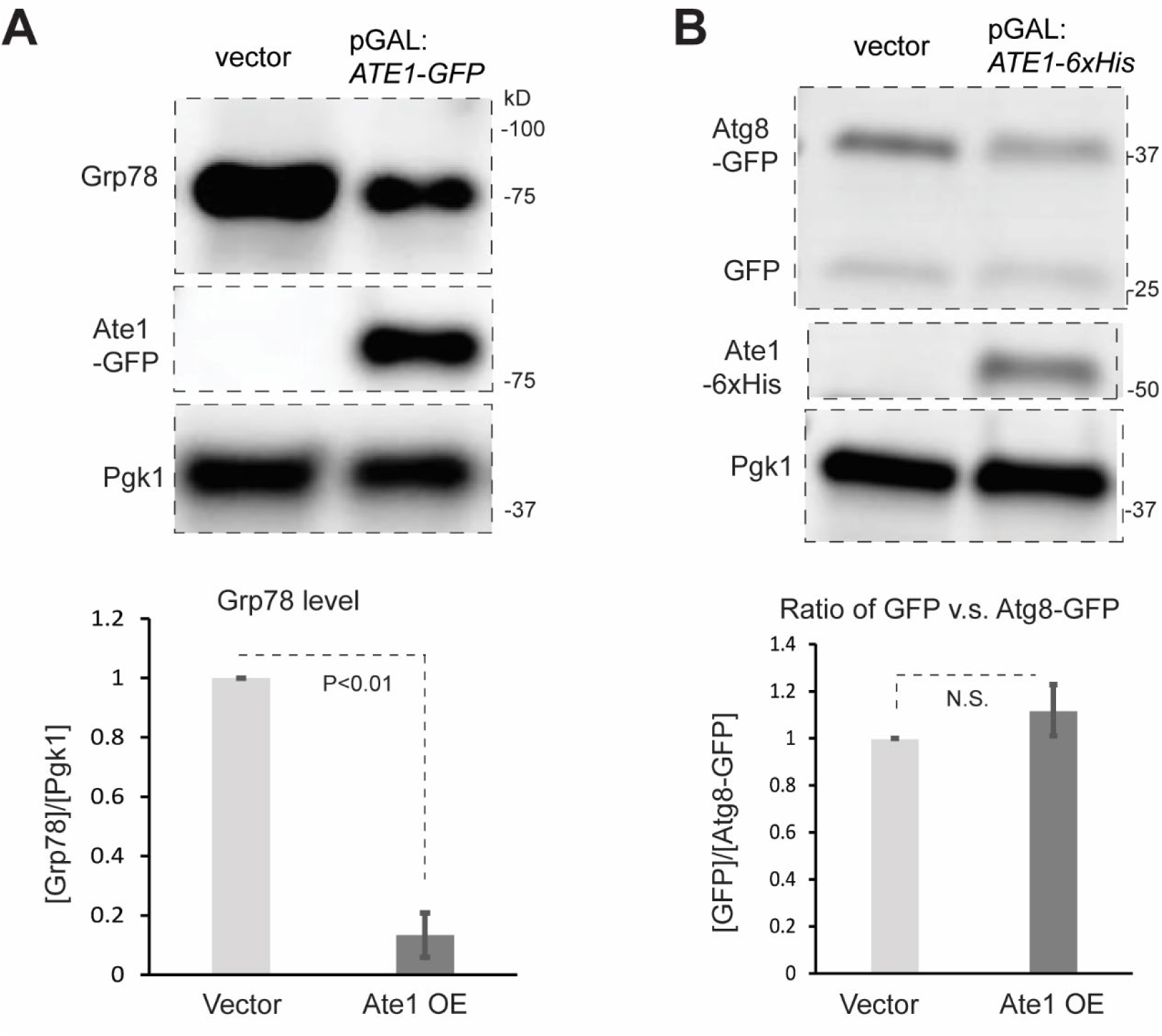
Ate1 overexpression does not significantly increase Endoplasmic Reticulum (ER) stress response or the autophagy degradation pathways. A) Top panel shows the representative Western blot for the level of Grp78/HDEL, a maker of the endoplasmic reticulum stress response, in yeast cells carrying the pYES2-pGAL:*ATE1-6xHis-*URA3 expression vector or the empty control vector, which were induced with 2% galactose for 6h in liquid media. Pgk1 is used as loading controls. Ate1 was probed with anti-GFP. Bottom panel is a graph indicating the fold differences in GRP78 levels between the Ate1 overexpression (OE) and vector control. Error bar denotes SEM (N=3). B) Top panel shows representative Western blot for the level of full-length Atg8-GFP and the derivative GFP, which is resulted form proteolysis in autophagy. The yeast cells carrying the pYES2-pGAL:*ATE1-6xHis-*URA3 expression vector or the empty control vector were induced with 2% galactose for 6h in liquid media. Pgk1 is a loading control. Bottom panel is a graph indicating the fold differences in the ratio between GFP and the full-length Atg8-GFP, which reflects the activity of autophagy, in the presence of Ate1 overexpression (OE) and vector control. Error bar denotes SEM (N=3).

## DISCUSSION

Ate1 (and with its gene duplicate ATE2 in plants) is the sole enzyme in most eukaryotes that catalyses arginylation, a PTM that has been compared to phosphorylation because they both add a charged and bulky chemical group to the target protein. While a substantial amount of data has unambiguously indicated the involvements of Ate1 or arginylation in cell death, the mechanism and even the exact role of Ate1/arginylation in such processes remain controversial due to conflicting reports. Our finding for the first time has pointed out that Ate1 regulates PCD mainly by driving the intrinsic, mitochondria-dependent apoptotic pathway.

Mitochondria play a central role in many PCD-related scenarios. These include but are not limited to cancer, embryogenesis, tissue development and regeneration, injuries, and aging. By demonstrating the role of Ate1 as a driver of the mitochondrial PCD pathway, our study thus provides important mechanistic insights for understanding the above-mentioned conditions, all of which are also known to involve Ate1 or arginylation. Furthermore, our finding shows that Ate1 is involved in the activation of mPTP and MOMP processes, which are also the most conserved mechanisms in PCD. This finding of Ate1 being involved in a fundamental cellular process is consistent to the nature of Ate1 as a gene transfer from α-proteobacteria --the ancestor of mitochondria. However, considering how many extensive studies have been invested in mPTP and MOMP, and considering the prominent effects of Ate1 in cell death observed by multiple groups, one may be surprised why the role of Ate1 as a basic regulator of a fundamental process remained overlooked for so long. Such a gap of knowledge further highlights the nature of the studies of Ate1/arginylation as a nascent research direction. This also indicates that the mechanisms governing mitochondrial PCD is likely more complex than previously assumed.

In our study, we showed that the elevation of Ate1 lead to the activation of mPTP and MOMP. We also showed that the deletion of the *AIF1*, one of the mitochondria-contained factors that can independently execute PCD, suppresses the Ate1-induced cell death. However, little or no effects were observed with the deletion of *NUC1* (yeast EndoG) (Fig.5D). Similarly, the deletion of yeast metacaspase (*YCA1*), which is expected to respond to the release of Cyc and trigger the execution of apoptosis, does not prevent Ate1-mediated lethality (Fig.5D). While it is still too early to make a conclusion at this stage, it is possible that AIF takes a more dominant role in the execution of yeast apoptotic program, while Cyc, caspase, or EndoG play redundant roles.

Contrast to the well-known role of Ate1 as a global regulator of cytosolic proteome, the PCD process induced by Ate1 elevation, as seen in response to acute stress and/or injury, appears to largely involve the mitochondrial pathway. However, we need to point out that our data did not rule out the potential involvements of cytosolic pathways. Instead, our data indicated several novel interactions that were not previously considered and should be further investigated to obtain a better understanding of PCD. For example, our data suggest that the elevated Ate1 appears to suppress the UPR signals (Fig.9A), which by itself can also trigger PCD given the right context. It is possible that Ate1 and UPR constitute negative feedback or a competing relationship for the regulation of PCD. While still speculative, this effect may be explained by the known impact of arginylation on the turnover of some of the pro-apoptotic proteins or protein fragments associated with UPR. These include calreticulin and Grp78, which may be retro-transported back to cytosol from ER during the crisis of UPR. Recent studies also showed that arginylation likely promotes autophagy[21, 22, 28]. However, in our tests the over-expressed Ate1 did not lead to significant changes of autophagy makers (Fig.9B). This discrepancy may be explained by considering multiple factors. For example, it is possible the effects of Ate1 in autophagy is only significant during proteolysis stress, which was not involved in the test condition in our study. Finally, Ate1/arginylation is conventionally considered a key player in the ubiquitination and proteasome-dependent degradation of many proteins in the cytosol. While this role does not appear to be directly required for the effects of Ate1 in the execution of the mitochondrial cell death pathways, our study was not designed to rule out its relevance for the upstream events leading to the elevation of Ate1 or the interactions of the cytosolic pathways. It is possible that the role of Ate1/arginylation in UPS is more complex in the context of cell death than previously assumed. A full understanding of these processes will be clarified, as these questions are investigated.

Our new findings also introduce new questions that were previously non-existent. Particularly, we are curious how Ate1 regulates mPTP and MOMP processes. Our current data appear to indicate that these are mainly controlled by the intra-mitochondrial Ate1. However, this question cannot be easily addressed right now because most existing studies about Ate1 were focused on the cytosol or nucleus. While a couple of mitochondrial proteins were sporadically identified in past screenings for arginylation substrates, most of them have no obvious relevancies to cell death. Further convoluting is that those studies were not even designed to include mitochondrial fraction in the first place. As such, it would be premature to speculate how Ate1 trigger the mitochondrial cell death pathway. This will have to wait for future investigations.

## MATERIALS AND METHODS

### Yeast strains

The *Saccharomyces cerevisiae* strains, wild-type or knockout, are mainly based on the parental strain BY4741 (*MATa his3Δ1 leu2Δ0 met15Δ0 ura3Δ0*) unless otherwise indicated. In several experiments where knockouts are not available for the BY4741 stain, the W303-1A strain background (*MATa leu2-3,112 trp1-1 can1-100 ura3-1 ade2-1 his3-11,15 ybp1-1*) was used. These yeasts were obtained from Horizon Discovery. The identity of each knockout strain was confirmed with PCR genotyping. The strain containing a GFP fused to the endogenous Ate1 was based on the BY4741 strain and was originally obtained from Open Biosystems (now part of Horizon Discovery) as described in our previous work[9].

### Culture of Yeast

The below yeast culture media were used:

YPD media: 2% glucose, 1% yeast extract, 2% peptone.

SD (Synthetic Defined) Media (per 1000ml): Yeast Nitrogen Base, 1.7g, Ammonium sulphate, 5 g, Dextrose/galactose/raffinose, 20g, required amino acids, 50mg, uracil (if required), 50mg.

To prepare solid media plates, 2% agar was added to the liquid media before autoclave.

The yeasts were incubated at 30°C to grow, unless otherwise indicated.

For the serial dilution growth assays, a single colony of yeast was inoculated in liquid medium and allowed to grow to the log phase. The spot platting of serial dilutions was performed as described before [52].

### Yeast cell transformation

Mid-log phase yeast cells with optical density (O.D.) at 600nm corresponding to 0.6 was grown in YPD liquid media and harvested. Yeast cells were washed twice with sterile water and added with transformation reaction mix containing polyethylene glycol (PEG 3000), lithium acetate (LiAce), carrier DNA (salmon sperm DNA) and desired plasmid. Cells were vortexed completely and subjected to heat shock at 42*C for 15 minutes as described earlier[9]. Transformants were selected on SD agar plate lacking essential supplements such as either uracil or leucine or tryptophan based on the experimental requirements.

### Molecular cloning and preparation of plasmids

High-fidelity DNA polymerases such as Herculase or Pfu turbo (from Agilent) were used to perform the DNA amplification. Restriction digestion and ligation were performed with corresponding restriction enzymes and T4-DNA ligase (NEB catalogue number M0202S) from New England Biolab. Chemical competent E.coli TOP10 (Thermo-Fisher Scientific) were used for plasmid transformation.

PCR reactions were performed in a Veriti Thermo Cycler (Applied Biosystems) or a T100 Thermo Cycler (BioRad). Most primers were ordered from Sigma-Aldrich.

The vector with the “pGAL” promoter in this manuscript is referred to the PYES2 plasmid, which contains the galactose driven promoter. It was obtained from (Life Technologies, # V825-20).

The construct (PYES2-P_GAL1_-Cyb2-Ate1-GFP) expressing mitochondrion localized (matrix) Ate1 was modified from the parental Ate1-GFP yeast expression plasmid was described in (1). A three-part ligation strategy was used. Fragment 1 was generated by double digestion of PYES2 vector with Age1 and Xba1 restriction enzyme. Fragment 2 harbouring the matrix targeting sequence of S.c. Cytochrome b2 (Cyb2) was PCR amplified from pYX233-b2delta-DHFR (addgene plasmid #163760) using the forward primer GTCCTCGTCTTCACCGGTCGCGTTCCTGA and the reverse primer GGGGTACCTTGAAGGGGACCCAATTTTTTCTCGGG. The stop-transfer signal (transmembrane domain) of cytochrome b2 was deleted (19 amino acids) so that the protein is targeted to the mitochondrial matrix (2). Fragment 2 was restriction digested using Age1 and Kpn1 enzymes. Fragment 3 containing Ate1-GFP was restriction digested and gel purified from PYES2-P_GAL1_-Ate1-GFP vector and subjected to three-part ligation using the T4 DNA ligase (NEB catalogue number M0202S). Culturing of yeast strains were carried out in YP media containing 2% glucose as a carbon source. Transformants of Wild-type BY4741 or W303A strains were generated by chemical transformation method (3) either using plasmid expressing Ate1-GFP, Ate1-6HIS or Cyb2-Ate1-GFP or its isogenic empty vector. Transformants were selected and grown either in synthetically defined agar or liquid media lacking uracil at 30°C.

The cloning of DD-β15-mCherryFP, arginylation Reporter: the cDNA of arginylation Reporter contains an N-terminal ubiquitin followed with a stretch of 15 amino acid from the N-terminus of beta actin (DDIAALVVDNGSGMC)[53], a linker region, and a then coding region for C-terminal mCherryFP. It was modified from a similar reporter DD-β15-GFP as described in our previous work[9]. The construct is cloned in a yeast expression vector, pGPD2 (Addgene # 43972), at EcoR1 and Xho1 sites. The following primers were used for this cloning: 5’–ATGTGCGGGATCCACCG–3’ and 5’-ACCTCGAGTTAATGGTGATGGT -3’.

The cloning of pGAL-Ate1-GFP-Ras2: It is modified from the original pGAL-Ate1-GFP, which was hosted at the pYES2.0 yeast expression backbone vector as described in our past work[9]. The sequence for the last 37 amino acids of Ras2 was amplified from the Ras2 gene contained in a PYES2 plasmid; its C-terminus was amplified using a Ras2-specific forward (ATATGCATGCAATAGTAAGGCCGGTCAAG) primer containing an SphI cutsite, and a plasmid-specific reverse primer (CGTACACGCGTCTGTAC). The amplified PCR product was extracted and digested with SphI and SalI before being ligated into a similarly digested Ate1-GFP-containing plasmid.

The inducible BAX-expression vector (pBM272-GAL-BAX-URA), with mouse BAX gene under Gal promoter and a URA3 selection marker, was acquired from Addgene (Plasmid #8770)

The constitutively expression vector for Bcl-xL (pBF339-ADH-BCLxL-TRP), with mammalian Bcl-XL gene under ADH1 promoter and a TRP1 selection marker, was acquired from Addgene (Plasmid #8773).

The constitutively expression vector for SOD1 and SOD2 are pGPD2-LEU2, which contains a GPD promoter and a LEU2 selection marker.

### Induction of Ate1 expression in yeast cells in liquid media

The yeasts (carrying Ate1 expression vectors with URA3 selection markers, or the empty vector control) were grown in SC -Ura liquid media supplemented with 2% raffinose until O.D. 600nm reach 0.5-0.6. Expression of Ate1 was achieved by adding 2% galactose at 30°C for 5-6 hours, which is not expected to trigger major cell death events by the overexpression itself [9].

To induce oxidative stress, cells were treated with 5% NaN_3_ for 30 minutes at 30°C.

### Chemical treatments of yeast cells on solid media plate

Yeast plates were made as previously described. To make the various inhibitors containing plates, a working stock was created using suitable solvent (DMSO for oligomycin/quercetin; ddH_2_0 for alpha-ketogluterate). After autoclaving the agar-containing media, it was allowed to cool till it was tolerable to touch (below 50°C). At this point the required amount of the chemical was added (from the working stock) into the media and allowed to stir till solution was homogeneous. Plates were then poured using this homogenous solution, and allowed to solidify for one day before use in experiment.

### Preparation of whole-cell lysates and mitochondrial fraction

To prepare whole-cell lysates, the yeast cells were lyzed by vortexing with glass beads in a 2x Laemmli SDS-loading buffer (for direct analysis). Alternatively, total protein extracts were prepared using sodium hydroxide and trichloroacetic acid as described previously[54].

To prepare mitochondrial fraction, the yeast cells were harvested by centrifugation. The mitochondria were isolated as per the protocol described by the manufacturer (abcam#ab178779) with following modifications. Briefly, yeast cells were harvested and rinsed with water. Cell pellets were either stored at -80°C or processed immediately. Cells were suspended in Buffer-A containing 1Mm DDT and incubated at 30°C with gentle shaking for 10 minutes. Cells were pelleted and resuspended with buffer-B containing lyticase. Cell suspension was incubated at 30°C for 30-45 minutes. The successful formation of spheroplast was validated under optical microscope. After centrifugation at 600xg for 5 minutes a lysis buffer containing protease inhibitor cocktail was added to the pellet and resuspended completely. Cell lysis was carried out using Dounce homogenizer 1X15 strokes on ice. Unlysed cell debris were pelleted by centrifugation at 800x g for 5 minutes twice. The supernatant containing mitochondria was pelleted by centrifugation at 12,000xg for 10 minutes. To remove contaminated cytosolic and nuclear proteins the resuspended mitochondrion fraction was treated with proteinase K at 5µg/ml concentration for 45 minutes on ice. The reaction was stopped by adding 20% TCA. Proteins were pelleted by centrifugation at 18,000xg for 15 minutes at 30°C. The protein pellet was washed once with ice-cold 100% acetone and centrifuged at 18,000xg.

For the cytochrome c-related fractionation, the following procedure was followed: Yeast cells were induced with galactose, harvested, and washed once with 1x PBS. Yeast cells were then resuspended in 650 mM DTT and 250 mM EDTA and incubated for 10 minutes at 30°C and 75-rpm gentle shaking. Cells were then spun down, supernatant removed, and washed with Sorbitol Buffer (1 M sorbitol, 0.15 K2HPO4, pH 7.4) once. Afterwards, the net weight of the yeast pellet was taken and yeast were resuspended in a mixture of sorbitol buffer (7 mL/g wet weight of pellet) and Zymolase (3 mg/g wet weight of pellet) and shaken at 50 rpm for 35 minutes at 30°C. After digestion with Zymolase, the resulting spheroplasts were pelleted at 3000x g and washed gently with sorbitol buffer. Pellets were then placed into a pre-chilled douncer homogenizer on ice and dounced a total of 10 times. A final volume of 100 uL of homogenate was taken (and if needed, sorbitol buffer was added) and spun down at 5000x g for 10 minutes at 4°C on a desktop centrifuge. Supernatant was isolated and placed in a separate tube on ice. This isolated supernatant was the centrifuged at 15000x g for 20 min at 4°C. The resulting supernatant was isolated from this and collected as the “cytosolic” fraction, while the residual pellet was considered the “mitochondrial” fraction. 2x Laemelli buffer was added to each fraction to each an equal final volume, and samples were boiled for 10 minutes before use in SDS-PAGE analysis.

### Analysis with SDS-PAGE and western blot

All protein samples were prepared in Laemmli SDS-loading buffer and heated in a boiling water bath for 10 minutes as described [53]. The samples were separated by electrophoresis on SDS-PAGE. Unless otherwise indicated, 4–20% Mini-PROTEAN® TGX™ Precast Protein Gels from Bio-Rad were used.

The freshly separated bands were transferred to nitrocellulose membrane for western blot analysis.

Western blot blocking reagent was obtained from Roche (catalogue number 75255200)

The primary antibodies include:

monoclonal mouse anti-GFP (from Roche, clone 7.1 and 13.1, Cat# 11814460001)

Rabbit anti-yeast alpha tubulin (Abcam EPR13799)

anti-yeast-phosphoglycerate kinase1 (Pgk1) (Thermofischer scientific # 459250),

Rabbit anti-yeast Cmc2 (gift from Dr. Antonio Barrientos)

mouse anti-Grp78 (SCBT# HDEL Antibody (2E7): sc-53472)

HSP70 Monoclonal antibody (Proteintech catalogue# 66183-1)

The custom-produced rabbit anti-RDD antibody was ordered from Genscript INC as described in our previous work [9].

The secondary antibodies include:

Goat anti-Mouse IgG (H+L) Secondary Antibody, HRP (Invitrogen catalogue# 314303)

Goat anti-Rabbit IgG (H+L) Secondary Antibody, HRP (Invitrogen catalogue# 656120

To visualize the antibody signal, the following chemiluminescent kits were used:

Pierce^TM^ ECL western catalogue # 32106

SuperSignal™ West Femto Maximum Sensitivity Substrate catalogue # 34096

The chemiluminescent signals were documented by a GE Amersham Imager model 600 and analysed with an ImageQuant TL software pack (v8.1) and its 1D gel analysis module.

### Microscopy

Optical and fluorescent imaging of the cells were done using a Zeiss Observer microscope, which was equipped with a series of objectives and the Zen-Pro analysis software.

### Flow cytometry (FACS)

FACS analysis or sorting was performed on a BD Canto-II flow cytometer, which is hosted by the flowcytometry core facility of the Sylvester Comprehensive Cancer Center at the University of Miami.

### Evaluation of yeast cell viability under Ate1 overexpression by serial dilution assay

The yeast cells were transformed with the overexpression vectors carrying different constructs of *ATE1*, in comparison of the empty vector pYES2, all of which contain the URA3 selection gene markers. The yeast cells were grown to mid-log phase in a synthetically defined media lacking uracil (sunrise science products; Catalog #: 1306-030). Optical density of yeast cells at 600nm corresponding to 0.5 was harvested and thoroughly washed with sterile water. Cells were serially diluted and spotted on to SD-URA plates containing 2% of glucose, raffinose, or galactose, unless otherwise indicated. Plates were incubated at 30°C for 2-5 days and photographed.

### TUNEL assay for apoptosis

Yeast cells were first grown in raffinose-containing media for 20 hours before being washed twice and resuspended in liquid dropout media containing 2% galactose (SDGal -Ura) and placed into an erlynmer flask (culture volume < 10% flask volume). Flasks were then placed into a shaking incubator (rpm 220) and incubated at 30°C for 6 hours. After, 1 O.D. worth of cells were harvested and washed once with galactose-containing dropout media and resuspended in 1 mL of this same media, in an Eppendorf tube. Formaldehyde was added to final concentration of 3.7% (v/v), and the tubes were mixed by inversion several times before being allowed to sit at room temperature for 30 minutes. Cells were then washed three times with PBS, before being digested with Zymolase 100T (24 ug/mL final; in PBS) at 37C for 60 minutes. Resulting spheroplasts were gently washed once with PBS before being applied to a microscope slide, and allowed to dry out (40 min at 37C). The rest of the procedure followed manufacturers instruction for “tissue section staining” using the Invitrogen Click-iT Plus TUNEL Assay.

### Statistics

Statistical significance was derived using a paired two-tailed student’s test.

## ACKNOWLEDGEMENTS

We thank the Sylvester Comprehensive Cancer Center (SCCC) at the University of Miami (UM) for providing access and subsidy for the flowcytometry core facilities.

The cost of experiments and the salary of Corin O’Shea, Ganapathi Kandasamy, Evan Ambrose and Fangliang Zhang are supported in part by National Institutes of Health (NIH) R01 grants GM138557.

Some of the experiments were performed from the financial support provided to Akhilesh Kumar by DST-SRG/2019/001360 and MoE-STARS/STARS1/385, India.

## Abbreviations

Ate1: arginyltransferase 1
GFP: green-fluorescence protein
KD: knockdown
KO: knock-out
WT: wild-type

## References

1. Kaji, H., G.D. Novelli, and A. Kaji, A Soluble Amino Acid-Incorporating System from Rat Liver. Biochim Biophys Acta, 1963. 76: p. 474–7.

2. Bachmair, A., D. Finley, and A. Varshavsky, In vivo half-life of a protein is a function of its amino-terminal residue. Science, 1986. 234(4773): p. 179-86.

3. Varshavsky, A., The N-end rule pathway and regulation by proteolysis. Protein Sci, 2011.

4. Chakraborty, G., et al., Posttranslational protein modification by amino acid addition in regenerating optic nerves of goldfish. J Neurochem, 1986. 46(3): p. 726–32.

5. Shyne-Athwal, S., et al., Protein modification by amino acid addition is increased in crushed sciatic but not optic nerves. Science, 1986. 231(4738): p. 603-5.

6. Shyne-Athwal, S., et al., Comparison of posttranslational protein modification by amino acid addition after crush injury to sciatic and optic nerves of rats. Exp Neurol, 1988. 99(2): p. 281–95.

7. Luo, D., G. Chakraborty, and N.A. Ingoglia, Post-translational modification of proteins by arginine and lysine following crush injury and during regeneration of rat sciatic nerves. Restor Neurol Neurosci, 1990. 2(2): p. 53–61.

8. Jack, D.L., G. Chakraborty, and N.A. Ingoglia, Ubiquitin is associated with aggregates of arginine modified proteins in injured nerves. Neuroreport, 1992. 3(1): p. 47–50.

9. Kumar, A., et al., Posttranslational arginylation enzyme Ate1 affects DNA mutagenesis by regulating stress response. Cell Death Dis, 2016. 7(9): p. e2378.

10. Masdehors, P., et al., Ubiquitin-dependent protein processing controls radiation-induced apoptosis through the N-end rule pathway. Exp Cell Res, 2000. 257(1): p. 48–57.

11. Birnbaum, M.D., et al., Reduced Arginyltransferase 1 is a driver and a potential prognostic indicator of prostate cancer metastasis. Oncogene, 2019. 38(6): p. 838–851.

12. Kwon, Y.T., et al., An essential role of N-terminal arginylation in cardiovascular development. Science, 2002. 297(5578): p. 96-9.

13. Brower, C.S. and A. Varshavsky, Ablation of arginylation in the mouse N-end rule pathway: loss of fat, higher metabolic rate, damaged spermatogenesis, and neurological perturbations. PLoS One, 2009. 4(11): p. e7757.

14. Kurosaka, S., et al., Arginylation regulates myofibrils to maintain heart function and prevent dilated cardiomyopathy. J Mol Cell Cardiol, 2012. 53(3): p. 333–41.

15. Saha, S. and A. Kashina, Posttranslational arginylation as a global biological regulator. Dev Biol, 2011. 358(1): p. 1–8.

16. Wang, J., et al., Arginyltransferase ATE1 is targeted to the neuronal growth cones and regulates neurite outgrowth during brain development. Dev Biol, 2017. 430(1): p. 41–51.

17. Rai, R., et al., Arginyltransferase suppresses cell tumorigenic potential and inversely correlates with metastases in human cancers. Oncogene, 2015.

18. Wang, Z., et al., A picorna-like virus suppresses the N-end rule pathway to inhibit apoptosis. Elife, 2017. 6.

19. Chui, A.J., et al., N-terminal degradation activates the NLRP1B inflammasome. Science, 2019. 364(6435): p. 82-85.

20. Wickliffe, K.E., S.H. Leppla, and M. Moayeri, Killing of macrophages by anthrax lethal toxin: involvement of the N-end rule pathway. Cell Microbiol, 2008. 10(6): p. 1352–62.

21. Cha-Molstad, H., et al., Amino-terminal arginylation targets endoplasmic reticulum chaperone BiP for autophagy through p62 binding. Nat Cell Biol, 2015. 17(7): p. 917–29.

22. Kim, H.J., et al., Crosstalk between HSPA5 arginylation and sequential ubiquitination leads to AKT degradation through autophagy flux. Autophagy, 2020: p. 1–19.

23. Decca, M.B., et al., Post-translational arginylation of calreticulin: a new isospecies of calreticulin component of stress granules. J Biol Chem, 2007. 282(11): p. 8237–45.

24. Brower, C.S., K.I. Piatkov, and A. Varshavsky, Neurodegeneration-associated protein fragments as short-lived substrates of the N-end rule pathway. Mol Cell, 2013. 50(2): p. 161–71.

25. Piatkov, K.I., C.S. Brower, and A. Varshavsky, The N-end rule pathway counteracts cell death by destroying proapoptotic protein fragments. Proc Natl Acad Sci U S A, 2012. 109(27): p. E1839–47.

26. Lopez Sambrooks, C., M.A. Carpio, and M.E. Hallak, Arginylated calreticulin at plasma membrane increases susceptibility of cells to apoptosis. J Biol Chem, 2012. 287(26): p. 22043–54.

27. Deka, K., et al., Protein arginylation regulates cellular stress response by stabilizing HSP70 and HSP40 transcripts. Cell Death Discov, 2016. 2: p. 16074.

28. Cha-Molstad, H., et al., Modulation of SQSTM1/p62 activity by N-terminal arginylation of the endoplasmic reticulum chaperone HSPA5/GRP78/BiP. Autophagy, 2016. 12(2): p. 426–8.

29. Beutner, G., et al., The Mitochondrial Permeability Transition Pore and ATP Synthase. Handb Exp Pharmacol, 2017. 240: p. 21–46.

30. Bock, F.J. and S.W.G. Tait, Mitochondria as multifaceted regulators of cell death. Nat Rev Mol Cell Biol, 2020. 21(2): p. 85–100.

31. Jiang, C., et al., Regulation of Mitochondrial Respiratory Chain Complex Levels, Organization, and Function by Arginyltransferase 1. Front Cell Dev Biol, 2020. 8: p. 603688.

32. Chen, L. and A. Kashina, Arginylation Regulates Cytoskeleton Organization and Cell Division and Affects Mitochondria in Fission Yeast. Mol Cell Biol, 2022. 42(11): p. e0026122.

33. Madeo, F., et al., Oxygen stress: a regulator of apoptosis in yeast. J Cell Biol, 1999. 145(4): p. 757–67.

34. Severin, F.F. and A.A. Hyman, Pheromone induces programmed cell death in S. cerevisiae. Curr Biol, 2002. 12(7): p. R233–5.

35. Herker, E., et al., Chronological aging leads to apoptosis in yeast. J Cell Biol, 2004. 164(4): p. 501–7.

36. Rai, R. and A. Kashina, Identification of mammalian arginyltransferases that modify a specific subset of protein substrates. Proc Natl Acad Sci U S A, 2005. 102(29): p. 10123–8.

37. Van, V., et al., Iron-sulfur clusters are involved in post-translational arginylation. Nat Commun, 2023. 14(1): p. 458.

38. Belotti, F., et al., Localization of Ras signaling complex in budding yeast. Biochim Biophys Acta, 2012. 1823(7): p. 1208–16.

39. Karakozova, M., et al., Arginylation of beta-actin regulates actin cytoskeleton and cell motility. Science, 2006. 313(5784): p. 192-6.

40. Kari, S., et al., Programmed cell death detection methods: a systematic review and a categorical comparison. Apoptosis, 2022. 27(7-8): p. 482–508.

41. Desagher, S. and J.C. Martinou, Mitochondria as the central control point of apoptosis. Trends in Cell Biology, 2000. 10(9): p. 369–377.

42. Wang, C. and R.J. Youle, The role of mitochondria in apoptosis*. Annu Rev Genet, 2009. 43: p. 95–118.

43. Gerle, C., Mitochondrial F-ATP synthase as the permeability transition pore. Pharmacol Res, 2020. 160: p. 105081.

44. Shchepina, L.A., et al., Oligomycin, inhibitor of the F0 part of H+-ATP-synthase, suppresses the TNF-induced apoptosis. Oncogene, 2002. 21(53): p. 8149–57.

45. Chin, R.M., et al., The metabolite alpha-ketoglutarate extends lifespan by inhibiting ATP synthase and TOR. Nature, 2014. 510(7505): p. 397-401.

46. Gledhill, J.R., et al., Mechanism of inhibition of bovine F1-ATPase by resveratrol and related polyphenols. Proc Natl Acad Sci U S A, 2007. 104(34): p. 13632–7.

47. Billen, L.P., et al., Bcl-XL inhibits membrane permeabilization by competing with Bax. PLoS Biol, 2008. 6(6): p. e147.

48. Rosa, N., et al., Bcl-xL acts as an inhibitor of IP3R channels, thereby antagonizing Ca2+-driven apoptosis. Cell Death and Differentiation, 2022. 29(4): p. 788–805.

49. Minn, A.J., et al., Bcl-xL regulates apoptosis by heterodimerization-dependent and - independent mechanisms. EMBO J, 1999. 18(3): p. 632–43.

50. Shimizu, S., Y. Shinohara, and Y. Tsujimoto, Bax and Bcl-x independently regulate apoptotic changes of yeast mitochondria that require VDAC but not adenine nucleotide translocator. Oncogene, 2000. 19(38): p. 4309–4318.

51. Moorthy, B.T., et al., The evolutionarily conserved arginyltransferase 1 mediates a pVHL-independent oxygen-sensing pathway in mammalian cells. Dev Cell, 2022. 57(5): p. 654–669 e9.

52. Kumar, A., et al., Mammalian proapoptotic factor ChaC1 and its homologues function as gamma-glutamyl cyclotransferases acting specifically on glutathione. EMBO Rep, 2012. 13(12): p. 1095–101.

53. Zhang, F., et al., Differential arginylation of actin isoforms is regulated by coding sequence-dependent degradation. Science, 2010. 329(5998): p. 1534-7.

54. Silve, S., et al., Membrane insertion of uracil permease, a polytopic yeast plasma membrane protein. Mol Cell Biol, 1991. 11(2): p. 1114–24.

